# The effects of age at menarche and first sexual intercourse on reproductive and behavioural outcomes: a Mendelian randomization study

**DOI:** 10.1101/423251

**Authors:** Rebecca B Lawn, Hannah M Sallis, Robyn E Wootton, Amy E Taylor, Perline Demange, Abigail Fraser, Ian S Penton-Voak, Marcus R Munafò

**Author notes:** Correspondence: Rebecca Lawn, School of Experimental Psychology, University of Bristol, 12a Priory Road, Bristol BS8 1TU, UK.; E.

## Abstract

There is substantial variation in the timing of significant reproductive life events such as menarche and first sexual intercourse. Life history theory explains this variation as an adaptive response to the developmental environment. In environments characterized by harsh conditions, adopting a fast life history strategy may increase fitness. In line with this, there is evidence demonstrating that greater childhood adversity is associated with earlier age at menarche. Here we applied Mendelian randomization (MR) methods to investigate whether there is a causal effect of variation in age at menarche and age at first sexual intercourse on outcomes related to reproduction, education and risky behaviour in UK Biobank (*N* = 114883–181,255). Our results suggest that earlier age at menarche affects some traits that characterize life history strategies including earlier age at first and last birth, decreased educational attainment, and decreased age at leaving education (for example, we found evidence for a 0.26 year decrease in age at first birth per year decrease in age at menarche, 95% confidence interval: −0.34 to −0.17; p < 0.0001). We find no clear evidence of effects of age at menarche on other outcomes, such as risk taking behaviour. Age at first sexual intercourse was also related to many life history outcomes, although there was evidence of horizontal pleiotropy which violates an assumption of MR and results should be treated with caution. Taken together, these results highlight how MR can be applied to test predictions of life history theory and to better understand determinants of health and social behaviour.

## Introduction

There is substantial variation between humans in the timing of significant reproductive life events such as age at menarche and first sexual intercourse (1,2). Life history theory explains this variation as an adaptive response to an individual’s developmental environment. Life history strategies are categorized as ‘fast’ (i.e., more effort directed towards reproduction such as earlier puberty and sexual activity) or ‘slow’ (i.e., later maturity and proportionally greater investment in a smaller number of offspring) (3,4). In some contexts, such as in developmental environments characterized by harsh conditions, adopting a fast life history strategy may increase reproductive fitness (1). Indeed, adverse childhood experiences have been shown to associate with earlier age at menarche, the start of a woman’s sexual maturity and reproductive potential (5). Age at first sexual intercourse is also related to the childhood environment (6–8). Whilst adopting a fast life history strategy may increase fitness, it may also have costs to the individual such as those associated with teenage pregnancy and risky behaviours such as violence, criminality, and substance abuse (3,9,10)). It is therefore important to examine how traits within life history strategies affect each other, especially when some of these traits may be modifiable.

Standard analytical approaches applied to observational data have been used to examine life history strategies in humans as it is not possible to manipulate developmental environments or reproductive timings (8,11). However, inferring causality in studies using such approaches is difficult, and likely to be affected by confounding (12). Mendelian randomization (MR) is an increasingly popular method in epidemiology for strengthening causal inference of risk factors that affect public health when randomized controlled trials and manipulation of risk factors is also not possible (13). It has been previously used to investigate psychological traits but has yet to be applied within an evolutionary framework (14). This method employs an instrumental variable analysis framework, with the instrument specifically being genetic variants known as single nucleotide polymorphisms (SNPs)(12). The instrument is used as a proxy for an exposure of interest (see Figure 1) (12,15,16).

**Figure 1.**
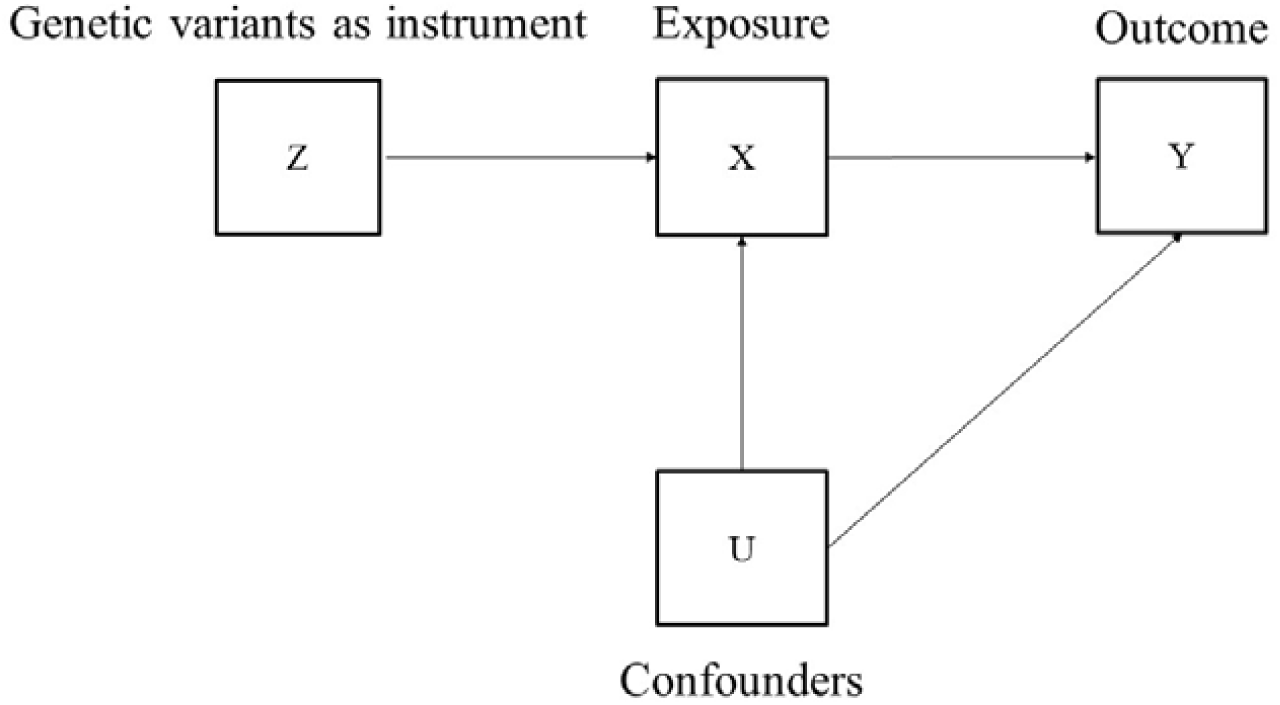
Diagram representing Mendelian randomization analysis.

Valid instrumental variables are defined by three main assumptions to allow for causal inference of results (15). First, the instrument is robustly associated with the exposure used in analysis (the relevance assumption). Genetic variants used as instrumental variables in MR are identified in genome-wide association studies (GWAS) to be significantly and independently associated with the exposure at a *p*-value less than 5×10^-8^. Second, the instrument is not associated with confounding factors (the independence assumption). By using genetic variants as instruments, MR exploits Mendel’s laws of segregation and independent assortment by which the inheritance of genetic variants is determined mostly independently of other genetic variants and the environment (12). This independence has been demonstrated through pairwise correlations between nongenetic variables and genetic variables, with genetic variants showing little association with each other (17). This highlights the advantages of using genetic variants as proxies of environmental exposures levels to overcome bias due to confounding to which non-genetic observational studies are prone (17). Third, the instrument only affects the outcome through the exposure (the exclusion restriction assumption). Violation of the exclusion restriction assumption where there is a direct effect of the genetic variant on the outcome not acting through the exposure is termed horizontal pleiotropy and is tested for in analyses (18,19). Additionally, since genotype is determined at conception, MR removes the risk of reverse causality (12,15). If these assumptions are met, effects estimated using MR should be free from bias due to confounding (12) and the associations between ZX and ZY (i.e., SNP-exposure and SNP-outcome) can be used to estimate the causal effect of X on Y(20). These SNP-exposure and SNP-outcome associations can be estimated from distinct non-overlapping samples of participants (20). One example of using MR to overcome biases when manipulation of the exposure is not practical is when investigating the effect of alcohol consumption on blood pressure and, ultimately, cardiovascular disease (previously described in (12) and (13)). Individuals who consume more alcohol may differ from individuals who consume less alcohol for other cardiovascular risk factors, such as by smoking more heavily. This could therefore introduce spurious associations due to bias from confounding by smoking heaviness. Using SNPs associated with metabolite responses to alcohol consumption is akin to randomizing individuals into higher or lower drinking conditions (20,21) and MR can therefore be used to estimate a causal effect. For more information on interpreting MR studies, see Davies et al. (13).

Here we apply the logic of MR to estimate the causal effects of one reproductive trait on reproductive and behavioural outcomes as a test of life history theory which postulates that early menarche and sexual intercourse (markers or results of exposure to ACEs) affect reproductive strategies to maximise fitness. Using instruments for age at menarche (and a separate instrument for age at first sexual intercourse), we independently investigate the effects of age at menarche and age at first sexual intercourse on several evolutionary relevant outcomes. A previous study (22) that included a sub-sample of participants from UK Biobank showed a causal effect of earlier age at menarche on earlier age at first birth, earlier age at last birth, earlier age at leaving education, increased alcohol intake, lower likelihood of being childless, greater number of children and decreased likelihood of remaining in education after 16 years. Additionally, earlier age at first sexual intercourse was causally related to earlier age at first birth, a greater number of children, increased likelihood of being an ever smoker, and decreased likelihood of attaining a degree. These findings suggest causal relationships between traits that characterize a life history strategy and support evolutionary explanations of variation in age at menarche and first sexual intercourse. We extend this work by using the full release of UK Biobank data (N = 114883–181,255) and a suite of novel methods to more robustly test for horizontal pleiotropy.

## Methods

### Exposure data

For our age at menarche instrument, we used independent SNPs associated with age at menarche (*p* < 5 × 10^-8^) from two GWAS (23,24). The first identified 123 SNPs (23) and explained approximately 3% of the observed variance in age at menarche (N = 182,416). The second identified 389 SNPs which explained about approximately 7% of the variance (N = 329,345). After excluding certain SNPs for methodological reasons (see Supplementary Text), we were left with 116 and 305 SNPs as instruments for age at menarche. Mean differences and standard errors (SE) for these SNPs and age at menarche associations in the GWAS discovery samples were recorded and this became our exposure data for age at menarche (see Supplementary Tables 1 and 2).

For our second instrument of age at first sexual intercourse, we used independent SNPs associated with age at first sexual intercourse (*p* < 5 × 10^-8^) (22) in both males and females. We recorded these GWAS associations, as done so for age at menarche, to be used as our exposure data for age at first sexual intercourse (see Supplementary Table 3). We used effect estimates identified in the pooled sex GWAS to increase statistical power. Our instrument consisted of 23 SNPs (see Supplementary Text for further details).

### Outcome data

The exposure associated SNPs described above were extracted from UK Biobank to derive SNP-outcome associations for each outcome. Extraction was done using PLINK (v2.00) and best guess algorithms for determining alleles.

#### Sample

UK Biobank is a population-based health research resource consisting of approximately 500 000 people, aged between 38 years and 73 years, who were recruited between the years 2006 and 2010 from across the UK (25). Particularly focused on identifying determinants of human diseases in middle-aged and older individuals, participants provided a range of information (including as demographics, health status, lifestyle measures, cognitive testing, personality self-report, and physical and mental health measures) via questionnaires and interviews (data available at www.ukbiobank.ac.uk). A full description of the study design, participants and quality control (QC) methods have been described in detail previously (26). UK Biobank received ethics approval from the Research Ethics Committee (REC reference for UK Biobank is 11/NW/0382). Genotyping information and quality checks in UK Biobank are described elsewhere (27).

#### Outcome measures

Our outcome measures were: age at first birth, age at last birth, reproductive period, number of children, childlessness, ever smoked, educational attainment in years, age when left education, alcohol intake, risk taking and number of sexual partners for those that indicated they had had sex. These measures were derived similarly to previous research (22,28). We re-coded data as missing if age at first sexual intercourse was younger than age at menarche; if age at leaving education was answered as having never attended school; at the 99.99^th^ percentile for number of children; at the 99.99^th^ percentile for number of sexual partners. Reproductive period was derived as the difference between age at last birth and age at first birth for those that had more than one child. To account for non-normal or categorical data, we included binary measures of childlessness (childlessness coded as 1), ever smoked (coded as 1 if participants had ever smoked in questions ‘Do you smoke tobacco now?’ or ‘In the past, how often have you smoked tobacco?’). Alcohol intake was a categorical variable indicating ‘never’ (coded as 6), ‘special occasions only’, ‘one to three times a month’, ‘once or twice a week’, three to four times a week’ and ‘daily or almost daily’ (coded as 1). Risk taking was measured as ‘yes’ (coded as 1) or ‘no’ responses to ‘Would you describe yourself as someone who takes risks?’. Only females were used for all outcome data.

### Data analysis

Data were harmonized to ensure that the effect of the SNP on the exposure and the SNP on the outcome corresponded to the same allele. The age increasing allele was used in order to conduct MR analyses and results were then reversed to report the effect of earlier age at menarche and first sexual intercourse. To derive the SNP-outcome associations for our outcome data, regressions were conducted in R adjusting for birth year and the top 10 genetic principal components. In sensitivity analysis, we additionally adjusted SNP-outcome associations for genotype array.

We used the 116 SNPs for age at menarche (23) in our main analysis as this GWAS did not include any UK Biobank data. For the 305 SNP instrument which includes some individuals from the UK Biobank (24), we calculated SNP-outcome associations and conducted analysis using outcome data from a UK Biobank sub-sample that did not overlap with the age at menarche GWAS. However, allocation into these sub-samples is related to smoking status (29) and division is therefore similar to stratifying on smoking. As smoking may be a collider in our analysis, this stratification could introduce bias. We therefore also derived SNP-outcome estimates and conducted analysis for the 305 SNP age at menarche instrument using the full UK Biobank sample, which will suffer from bias towards the observational estimate due to sample overlap with the GWAS of the exposure (30). It is also not possible to assess the suitability of one MR method (MR-Egger) with sample overlap as the suitability value (the I^2^_GX_ value) cannot be reliably measured. As the age at first sexual intercourse GWAS (22) was conducted solely in a sub-sample of UK Biobank participants, we conducted a fixed-effects meta-analysis of the SNP-outcome estimates in the full UK Biobank sample in addition to MR analysis in the non-overlapping sub-sample of UK Biobank. This fixed-effects meta-analysis is equivalent to performing an unweighted allele score analysis (31) and suffers from less bias than a weighted analysis.

Exposure and outcome data (i.e., SNP-exposure and SNP-outcome associations) were combined using multiple MR approaches (inverse variance weighted, weighted median, mode-based estimator and MR-Egger methods). These methods each rely on different assumptions regarding horizontal pleiotropy so that a consistent effect across all methods increases our confidence in results, although some methods suffer from reduced statistical power (19). An inverse variance weighted approach which is analogous to a weighted regression of SNP-outcome coefficients on SNP-exposure coefficients with the intercept constrained to zero (32,33), and further assumes all variants are valid instruments (i.e. meet the relevance, independence and exclusion restriction assumptions defined above) or allows pleiotropy to be balanced across instruments when using the random effects model (33). The weighted median estimates a causal effect if at least 50% of the data for analysis is from variants that are valid instruments (i.e., meet the relevance, independence and exclusion restriction assumptions defined above) (15,34). The mode-based estimator provides a causal estimate when the largest number of similar individual-instrument estimates come from valid instruments, even if the majority are invalid (35). A tuning parameter of 0.5 was set for mode-based estimator analysis. MR-Egger regression analysis, which does not constrain the intercept to zero and the intercept term therefore estimates overall horizontal pleiotropy was also conducted (19). In addition to these analyses, we conducted Radial MR and a leave one out analysis for age at first sexual intercourse which helps to identify outlier SNPs (36). For binary outcomes, all MR results were transformed to odds ratios by exponentiating them.

We calculated Cochran’s Q for these inverse variance weighted analyses to test if effects differ across variants (13). We further calculated the I^2^_GX_ to assess the suitability of MR-Egger where above 0.9 is desired (37), and mean F statistics which indicate the strength of the instrument. For age at first sexual intercourse, the unweighted I^2^_GX_ was low and we therefore performed a SIMEX adjustment with unweighted analysis.

Age at menarche analysis using the 116 SNP instrument was repeated after removing SNPs associated with body mass index at *p* < 5 × 10^-8^ (31,38,39). This resulted in 9 SNPs being removed (rs10938397, rs12446632, rs2947411, rs3101336, rs543874, rs7103411, rs7138803, rs7514705, rs8050136).

## Results

Mean age in our sample was 57 years (standard deviation [SD] = 7.91). Mean age at menarche and first sexual intercourse were 13 years (SD = 1.60) and 19 years (SD = 3.44), respectively. Further sample characteristics are given in Table 1. Further details of the instruments are provided in Supplementary Table 4.

**Table 1.** Population characteristics of UK Biobank sample used as outcome data.

|  | **Total N** | **Mean (SD) or N (%)** |
| --- | --- | --- |
| Age at assessment, years | 181 358 | 56.67 (7.91) |
| Age at menarche, years | 176 262 | 12.95 (1.60) |
| Age at first sex, years | 154 599 | 19.02 (3.44) |
| Age first birth, years | 124 093 | 25.39 (4.54) |
| Age last birth, years | 123 926 | 30.15 (4.80) |
| Reproductive period, years | 123 892 | 4.76 (3.65) |
| Number of sexual partners | 149 902 | 4.63 (6.99) |
| Number of children | 181 247 | 1.81 (1.15) |
| Childlessness | 181 255 |  |
| Yes |  | 33 242 (18.34) |
| No |  | 148 013 (81.66) |
| Age when left education, years | 124 279 | 16.63 (2.03) |
| Educational attainment, years | 179 731 | 13.05 (4.32) |
| Alcohol intake | 181 233 |  |
| Daily or almost daily |  | 30 918 (17.06) |
| Three or four times a week |  | 39 346 (21.71) |
| Once or twice a week |  | 47 864 (26.41) |
| One to three times a month |  | 23 723 (13.09) |
| Special occasions only |  | 25 101 (13.85) |
| Never |  | 14 281 (7.88) |
| Ever smoked | 180 751 |  |
| Yes |  | 101 112 (55.94) |
| No |  | 79 639 (44.06) |
| Risk taking | 174 718 |  |
| Yes |  | 31 973 (18.30) |
| No |  | 142 745 (81.70) |

### Age at menarche

Using the 116 SNP instrument for age at menarche we find consistent evidence of a causal effect of earlier age at menarche on earlier age at first birth across all MR methods. We find some evidence of an effect of earlier age at menarche on earlier age at last birth and all MR methods showed point estimates in a consistent direction. There was no clear evidence of an effect of age at menarche on duration of reproductive years, number of children, or number of sexual partners, and little evidence for an effect on likelihood of being childless with results showing confidence intervals consistent with the null and inconsistency for the direction of point estimates across MR approaches. These results are presented in Table 2 and 3.

**Table 2.** Estimates of the causal effect of earlier age at menarche (116 SNP instrument) on life history outcomes using full UK Biobank data.

|  | N for outcome | Inverse variance weighted |  | MR-Egger regression |  | Weighted median |  | Weighted mode-based estimator |  |
| --- | --- | --- | --- | --- | --- | --- | --- | --- | --- |
|  |  | Estimate (LCI, UCI) | P | Estimate (LCI, UCI) | P | Estimate (LCI, UCI) | P | Estimate (LCI, UCI) | P |
| Reproduction |  |  |  |  |  |  |  |  |  |
| Age first birth | 115070 - | -0.256 | <0.0001 | -0.260 | 0.051 | -0.325 | <0.0001 | -0.362 | 0.051 |
|  | 124093 | (-0.342, -0.171) |  | (-0.522, 0.002) |  | (-0.471, -0.178) |  | (-0.722, -0.002) |  |
| Age last birth | 114916 - | -0.235 | <0.0001 | -0.208 | 0.141 | -0.241 | 0.002 | -0.225 | 0.132 |
|  | 123926 | (-0.325, -0.144) |  | (-0.487, 0.07) |  | (-0.391, -0.091) |  | (-0.515, 0.066) |  |
| Reproductive period | 114883 - | 0.018 | 0.622 | 0.040 | 0.711 | 0.015 | 0.783 | -0.076 | 0.569 |
|  | 123892 | (-0.053, 0.088) |  | (-0.175, 0.255) |  | (-0.09, 0.12) |  | (-0.335, 0.184) |  |
| Number of sexual partners | 138920 - | -0.052 | 0.387 | 0.094 | 0.609 | -0.030 | 0.773 | -0.010 | 0.962 |
|  | 149902 | (-0.171, 0.067) |  | (-0.27, 0.459) |  | (-0.235, 0.175) |  | (-0.423, 0.403) |  |
| Number of children | 168050 - | -0.016 | 0.087 | 0.023 | 0.423 | -0.022 | 0.142 | 0.020 | 0.532 |
|  | 181247 | (-0.034, 0.002) |  | (-0.033, 0.078) |  | (-0.052, 0.007) |  | (-0.042, 0.082) |  |
| Childlessness | 168058 - | 1.060 | 0.006 | 0.987 | 0.838 | 1.047 | 0.184 | 1.038 | 0.592 |
|  | 181255 | (1.017, 1.105) |  | (0.869, 1.121) |  | (0.978, 1.121) |  | (0.905, 1.192) |  |
| Education |  |  |  |  |  |  |  |  |  |
| Age when left education | 115204 - | -0.062 | 0.002 | -0.153 | 0.011 | -0.095 | 0.004 | -0.126 | 0.119 |
|  | 124279 | (-0.100, -0.024) |  | (-0.27, -0.036) |  | (-0.158, -0.031) |  | (-0.283, 0.031) |  |
| Educational attainment in years | 166640 - | -0.072 | 0.036 | -0.237 | 0.025 | -0.128 | 0.047 | -0.246 | 0.047 |
|  | 179731 | (-0.139, -0.005) |  | (-0.443, -0.03) |  | (-0.253, -0.003) |  | (-0.487, -0.005) |  |
| Risky behaviours |  |  |  |  |  |  |  |  |  |
| Alcohol intake | 168039 - | 0.059 | <0.0001 | -0.030 | 0.427 | 0.035 | 0.102 | 0.007 | 0.863 |
|  | 181233 | (0.035, 0.083) |  | (-0.103, 0.044) |  | (-0.007, 0.077) |  | (-0.075, 0.089) |  |
| Ever smoked | 167584 -<br>180751 | 1.002<br>(0.970, 1.034) | 0.915 | 1.015<br>(0.921, 1.121) | 0.755 | 0.994<br>(0.942, 1.049) | 0.831 | 1.035<br>(0.924, 1.16) | 0.546 |
| Risk taking | 161994 -<br>174718 | 0.989<br>(0.949, 1.032) | 0.606 | 1.092<br>(0.96, 1.242) | 0.180 | 0.984<br>(0.916, 1.058) | 0.674 | 0.979<br>(0.832, 1.153) | 0.807 |

Note: Mendelian Randomization approaches used: inverse variance weighted, weighted mode-based estimator, MR-Egger regression and weighted median. (LCI: lower 95% confidence interval; UCI: upper 95% confidence interval).

**Table 3.**
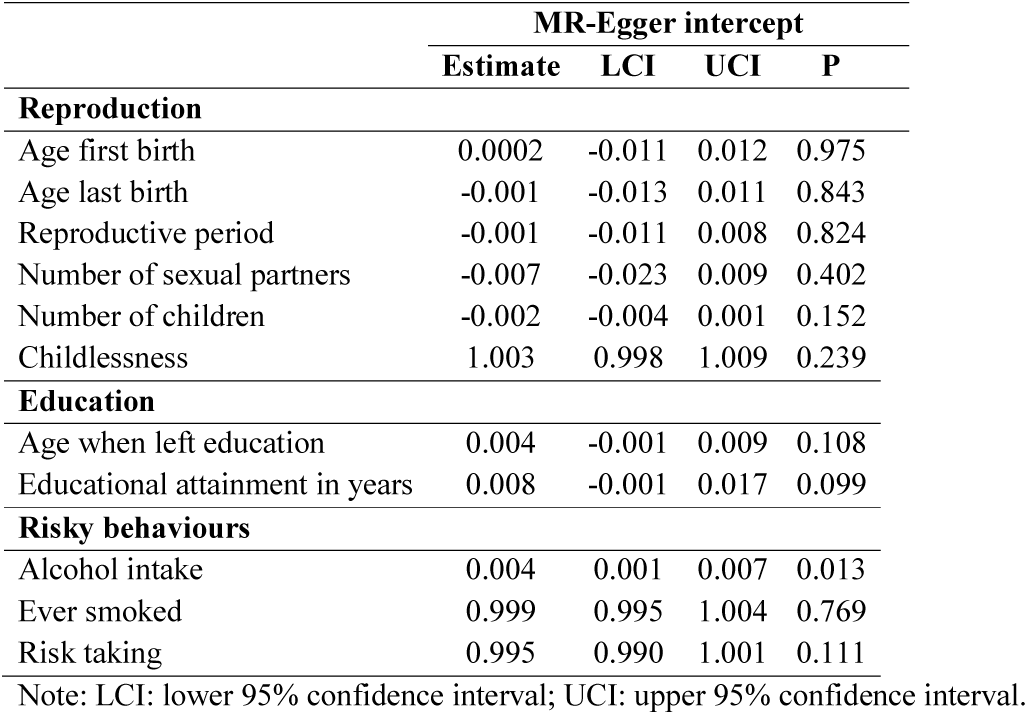
MR-Egger intercept values for age at menarche (116 SNP instrument) on life history outcomes using full UK Biobank data.

For education outcomes, there was evidence of an effect of earlier age at menarche on lower educational attainment and age at leaving education across most MR methods and consistent point estimates for all MR approaches (Table 2 and 3). Alcohol intake appeared to decrease with earlier age at menarche, but the MR-Egger intercept (p = 0.013) indicated horizontal pleiotropy, suggesting that this effect does not remain when horizontal pleiotropy is accounted for (Table 2 and 3). No clear evidence was found for effects of age at menarche on having ever smoked or risk taking behaviour although these measures were binary and therefore we had less statistical power to detect effects on these (Table 2).

After removing SNPs also associated with body mass index (31,39) from our genetic risk score, results were broadly similar to the main analysis although MR-Egger regression analysis showed decreased estimates and for many outcomes the p-values increased. This could be due to eliminating a possible pathway via body mass index and/or reduced statistical power associated with using fewer SNPs (Supplementary Tables 5 and 6). We repeated analyses using the 305 SNP instrument for age at menarche. Results were with broadly similar to the main analysis (Supplementary Tables 7-10). There was slight increased evidence for an effect on number of sexual partners, ever smoked and childlessness. This analysis suffers from greater bias as it is uses a sub-sample of UK Biobank (described above) or alternatively, when using the entire UK Biobank sample in analyses, it results in overlap between the exposure and outcome datasets, which has shown to bias results towards the observational estimate (30).

### Age at first sexual intercourse

We conducted a fixed effects meta-analysis of the 23 SNP-outcome associations in UK Biobank and found evidence of relationships for earlier age at first sexual intercourse with earlier age at first birth, earlier age at last birth, a longer reproductive period, increased number of sexual partners, a greater number of children, decreased likelihood of being childlessness, earlier age at leaving education, lower educational attainment, increased likelihood of having ever smoked and increased likelihood of risk taking behaviour (Table 4).

**Table 4.** Fixed effects meta-analysis of SNP-outcome associations using full UK Biobank and SNPs identified for age at first sexual intercourse (23 SNPs).

|  | Fixed effects meta-analysis |  |  |  |  |
| --- | --- | --- | --- | --- | --- |
|  | N | Estimate | LCI | UCI | P |
| **Reproduction** |  |  |  |  |  |
| Age first birth | 109021 - 124093 | -0.061 | -0.069 | -0.053 | <0.0001 |
| Age last birth | 108873 - 123926 | -0.046 | -0.055 | -0.037 | <0.0001 |
| Reproductive period | 108842 - 123892 | 0.015 | 0.008 | 0.022 | <0.0001 |
| Number of sexual partners | 131643 - 149902 | 0.019 | 0.007 | 0.030 | 0.002 |
| Number of children | 159140 - 181247 | 0.006 | 0.004 | 0.008 | <0.0001 |
| Childlessness | 159147 - 181255 | 0.986 | 0.982 | 0.990 | <0.0001 |
| **Education** |  |  |  |  |  |
| Age when left education | 109137 - 124279 | -0.011 | -0.015 | -0.007 | <0.0001 |
| Educational attainment in years | 157817 - 179731 | -0.015 | -0.022 | -0.008 | <0.0001 |
| **Risky behaviours** |  |  |  |  |  |
| Alcohol intake | 159137 - 181233 | 0.001 | -0.001 | 0.003 | 0.409 |
| Ever smoked | 158702 - 180751 | 1.010 | 1.007 | 1.014 | <0.0001 |
| Risk taking | 153432 - 174718 | 1.011 | 1.006 | 1.015 | <0.0001 |

Note: LCI: lower 95% confidence interval; UCI: upper 95% confidence interval.

We also used MR to examine effects between age at first sexual intercourse and these life history outcomes, taking SNP-exposure associations from a GWAS (22) and SNP-outcome associations in a sub-sample of UK Biobank, therefore likely affected by selection bias. There appeared to be a consistent effect of earlier age at first sexual intercourse on earlier age at first birth and earlier age at last birth and increased likelihood of risk taking behaviour across MR methods (Supplementary Tables 11). However, the MR-Egger intercept showed evidence of horizontal pleiotropy for most outcomes (Supplementary Tables 12). Results for Radial MR and a leave one out analysis suggested no strong influence of outliers (further details are provided in Supplementary Text).

## Discussion

The results suggest that earlier age at menarche is causally related to some traits that characterize a fast life history strategy, such as earlier age at first birth, earlier age at last birth, lower educational attainment, and earlier age at leaving education. This is consistent with previous findings (22,40). We also found no clear effect of age at menarche on number of children in this (female only) sample, supporting previous findings (22). Here, applying additional MR methods to those used previously, we find that the effect of age at menarche on alcohol intake is not robust (22).

We find mixed evidence for age at first sexual intercourse on these life history traits, with results suggesting possible violation of the exclusion restriction assumption of no direct effects of the instrument on the outcome not acting through the exposure (i.e., the presence of horizontal pleiotropy) (12,19). We detect the presence of horizontal pleiotropy on multiple outcomes, suggesting that previous findings may have also included pleiotropic effects (22). Results for age at first sexual intercourse are therefore not robust and the ability for casual inference is weakened.

The effects of earlier age at menarche on these reproductive and educational traits can be viewed as directing effort towards short-term reproductive goals and risky behaviour as an important part of a fast life history strategy (3). Variation in age at menarche may therefore represent an important causal component of a suite of adaptations (1). Earlier age at first child as part of a fast life history strategy can be considered an adaptive response to ACEs and our finding of an effect of earlier age at menarche on earlier age at first birth is therefore in line with this (41). It is however interesting that we see an effect of earlier age at menarche on earlier age at last birth, with no clear effect on reproductive period. This suggests that individuals on a fast life history strategy are not just starting their reproductive life earlier but shifting their reproductive life forward in time. Nettle highlights that individuals in more deprived areas with short life expectancy, likely on a fast life history strategy, need to reproduce earlier than individuals in more affluent areas with higher life expectancy to be in good health for an equivalent period of care (42). Education is a key predictor of positive later life outcomes in the UK, and our finding of a causal effect of earlier age at menarche on increased educational attainment provides important information for determinants of educational attainment which should be independent from confounding (40). Investing in education can be seen as a slow life history trait with delayed benefits (43). The effect of age at menarche on educational attainment may be due to variation in cognition following variation in age at menarche and gonadal hormones, due to menarche, that may influence behaviour during schooling (40,44).

As a component of life history strategy, we would have expected to see an effect of earlier age at menarche on increased number of children or likelihood of remaining childless, although access to contraception may influence this relationship. Number of sexual partners may not be as influenced by contraception as therefore can be used as a proxy for reproductive success (45). We did not find a clear effect of age at menarche on number of sexual partners although it may be due to female reproductive success not being as restrained by number of sexual partners as is males. It is further possible that the effect of menarche on number of children is masked by the detrimental effects of risky behaviours, such as substance use, on fertility in the modern environment (46,47). Although our results show no clear evidence of an effect of earlier age at menarche on increased risky behaviours and substance use, binary measures of smoking and risk taking were used, resulting in less statistical power. Furthermore, the measure of risk taking was a single item asking whether participants would describe themselves as someone who takes risks and may not capture the full extent of risk taking behaviour. We did not show an effect of age at menarche on alcohol intake, another form of substance use which has also been shown to be associated with decreased fertility (46,47). Further research should examine the mediating causal relationships between age at menarche and fertility in the modern environment using more detailed measures of substance use and larger samples.

Our study highlights how MR can be applied to test predictions within life history theory to provide evidence of causality and increase our understanding of health and social behaviour. A strength of the present study is the use of multiple MR methods. This allowed us to extend upon the findings of previous research (22) and triangulate across methods, each with varying and orthogonal assumptions, to provide greater confidence in results (48). We were further able to compare evidence using two instruments for age at menarche.

Additionally, we used a large population-based sample for our analysis to help identify the small effects common in genetic studies (32), although we acknowledge that for binary outcomes power was more limited. There are also a number of limitations to consider. Firstly, the age at first sexual intercourse GWAS was conducted in a sub-sample of UK Biobank data and we therefore conducted an unweighted analysis due to this sample overlap, using a fixed effects meta-analysis method. We additionally conducted MR by dividing our outcome sample to avoid overlap of participants, however this may have introduced bias as sub-division is related to smoking status and therefore akin to stratifying on smoking, which may be effected by our exposure and outcome (termed collider bias) (29). Secondly, we used SNPs for age at first sexual intercourse, and their associations, identified in pooled sex GWAS, due to reductions in power of using SNPs identified in females only and our exposure and outcome data therefore consists of different populations (20). Thirdly, the SNP-exposure associations were used from discovery analysis, which may cause upward bias of estimates (31,49), however using discovery data is common in MR studies and our unweighted analysis for age at first sexual intercourse did not use GWAS estimates (31,50). Fourthly, UK Biobank data is unrepresentative of the population, with a 5% response rate, and therefore and suffers from selection bias which may generate spurious associations (25,51). Finally, variants are non-specific and we cannot fully remove population structure, which can induce spurious associations through confounding, even within a sample of European ancestry and adjusting for principal components as done so here (52).

## Conclusions

We found some evidence that age at menarche is causally related to other life history traits and outcomes. Age at first sexual intercourse was also related to many life history outcomes, although there was evidence of horizontal pleiotropy which violates the exclusion restriction assumption of MR and results should therefore be treated with caution (18,19). This study highlights how analysis from genetic epidemiology can be used to answer how life history traits are related within life history strategies. There are increasing GWAS conducted on evolutionary relevant traits and future research could apply these MR techniques to further test predictions of life history theory, such as whether age at menarche is a mechanism between early life adversity and these evolutionary outcomes (53).

## Acknowledgments

RL, AF, REW, HMS, AET, and MRM work in the Medical Research Council Integrative Epidemiology Unit at the University of Bristol, which is supported by the Medical Research Council and the University of Bristol and funds RL’s PhD studentship (grant number: MC_UU_00011/7; MC_UU_12013/5). AF is supported by Career Development Awards from the UK Medical Research Council (MR/M009351/1). MRM is a member of the UK Centre for Tobacco and Alcohol Studies, a UKCRC Public Health Research: Centre of Excellence. This study was supported by the NIHR Biomedical Research Centre at the University Hospitals Bristol NHS Foundation Trust and the University of Bristol. The views expressed in this publication are those of the author(s) and not necessarily those of the NHS, the National Institute for Health Research or the Department of Health. Funding from British Heart Foundation, Cancer Research UK, Economic and Social Research Council, Medical Research Council, and the National Institute for Health Research, under the auspices of the UK Clinical Research Collaboration, is gratefully acknowledged. This research has been conducted using the UK Biobank Resource under Application number 6326. The authors would like to thank Dr Jack Bowden and Mr Wes Spiller for their technical assistance and Dr Ian Stephen for his helpful comments. The authors also thank Dr Ruth Mitchell, Dr Gibran Hemani, Mr Tom Dudding and Dr Lavinia Paternoster for conducting the quality control filtering of UK Biobank data.

## Author contributions

Contributors AF, MRM, ISP-V and RBL conceived the study. RBL conducted the analysis and drafted the initial manuscript. HS, REW, AET, and PD assisted with analysis. All authors assisted with interpretation, commented on drafts of the manuscript and approved the final version.

## Data accessibility

Genome wide summary data for age at menarche and age at first sexual intercourse can be downloaded from the ReproGen Consortium website (www.reprogen.org). Data for age at first sexual intercourse can be taken from the associated reference (www.nature.com/articles/ng.3551#references). UK Biobank data is available upon application (www.ukbiobank.ac.uk). Analysis scripts are available on GitHub (https://github.com/MRCIEU/AFS_AAM_LifeHistory.git).

